# Neutrophil extracellular trap stabilization leads to improved outcomes in murine models of sepsis

**DOI:** 10.1101/630483

**Authors:** Kandace Gollomp, Amrita Sarkar, Steven H Seeholzer, Lubica Rauova, M. Anna Kowalska, Mortimer Poncz

## Abstract

Sepsis is characterized by multi-organ system dysfunction that occurs due to infection. It is associated with unacceptably high morbidity and mortality and in need of improved therapeutic intervention. Neutrophils play a crucial role in sepsis, releasing neutrophil extracellular traps (NETs) composed of DNA complexed with histones and toxic antimicrobial proteins that ensnare pathogens but also damage host tissues. At presentation, patients likely have a significant NET burden contributing to the multi-organ damage. Therefore, interventions that prevent NET release would likely be ineffective at preventing NET-based injury. Treatments that enhance NET degradation may liberate captured bacteria and toxic NET degradation products (NDPs) and therefore be of limited therapeutic benefit. We propose that interventions that stabilize NETs and sequester NDPs may be protective in sepsis. Platelet factor 4 (PF4, CXCL4) a platelet-associated chemokine, binds and compacts NETs, increasing their resistance to deoxyribonuclease I. A monoclonal antibody, KKO, which binds to PF4-NET complexes, further enhances this resistance. We now show that PF4 increases NET-mediated bacterial capture in vitro, reduces the release of NDPs, and improves outcome in murine models of sepsis. An Fc-modified KKO further enhances deoxyribonuclease resistance, decreases NDP release, and increases survival in these models, supporting a novel NET-targeting approach to improve outcomes in sepsis.

## Introduction

Sepsis is defined as a dysregulated response to infection that leads to life-threatening organ damage. It affects millions of people each year and remains one of the most common causes of mortality worldwide^1,2^. Care consists of antibiotics and supportive measures^3^, but these interventions do not target the host response that causes much of the morbidity in sepsis^4^, and the poor survival rate has not changed significantly for several decades^5^. New emphasis is being placed on the identification of novel targeted interventions^6^ and the innate immune response has become an area of great interest^7^. Neutrophils, the most abundant circulating white blood cell^8^, play a crucial role in sepsis. They are recruited to inflamed vessels where they release neutrophil extracellular traps (NETs), webs of negatively charged cell-free DNA (cfDNA) complexed with positively-charged histones and antimicrobial proteins, such as myeloperoxidase (MPO) and neutrophil elastase (NE) that ensnare pathogens, limit bacterial spread^9^, and degrade inflammatory cytokines^10,11^. However, these positive effects occur at the expense of collateral tissue damage. NETs are lysed by circulating deoxyribonucleases (DNases)^12^, releasing NET degradation products (NDPs, e.g., cfDNA, cfDNA-MPO complexes, and histones) that exert harmful effects. For example, cfDNA triggers the contact-pathway^13^ and fixes complement^14^, MPO induces oxidative tissue damage^15^, NE cleaves tissue factor pathway inhibitor (TFPI) promoting thrombosis^16^, while histones cause platelet activation^17^ and are directly toxic to endothelial cells^18^. In septic patients, plasma levels of NDPs correlate with end-organ damage and mortality^19,20^. It has been suggested that preventing NET release (NETosis) might be beneficial^21^, but we believe that this strategy may be ineffective as septic patients likely have a large amount of NETs released by the time they become clinically ill. The use of DNase I to accelerate NET degradation has also been proposed as a therapeutic intervention^22^. However, results of prior studies with this intervention have been mixed^23^, raising the concern that DNase I treatment may lead to higher levels of circulating bacteria and the systemic release of NDPs^9,10,24^.

We propose an alternative strategy of NET-directed therapy in sepsis, in which NETs are compacted and stabilized, leading to NDP sequestration. Platelet factor 4 (PF4, CXCL4), a positively-charged chemokine released in high concentrations by activated platelets^25^, may exert this effect. PF4 binds to and aggregates polyanions like heparin^26^. We and others have found that PF4 similarly aggregates NETs, physically compacting them, and enhancing their resistance to endogenous and microbial nucleases^27,28^. We speculated that this activity may be, in part, responsible for our previous observation that PF4 enhances survival in murine lipopolysaccharide (LPS) endotoxemia^29^. KKO is a monoclonal antibody directed against PF4-heparin complexes that induces thrombocytopenia and a prothrombotic state when injected into mice that express human (h) PF4 and FcγRIIA, mimicking the clinical disorder of heparin-induced thrombocytopenia (HIT)^30^. KKO binds hPF4-NET complexes, further enhancing DNase resistance^27^. We speculated that treatment with an Fc-modified KKO that could not activate platelets via FcγRIIa may augment the protective effect of PF4 in sepsis by stabilizing hPF4-NET complexes. To investigate this hypothesis, we generated a deglycosylated version of KKO, termed DG-KKO, that retains the ability to bind hPF4-NET complexes, but has a reduced capacity to stimulate the inflammatory response. In mice treated with LPS and in the murine cecal ligation and puncture (CLP) model of polymicrobial sepsis, treatment with DG-KKO led to protection from thrombocytopenia, decreased plasma NDP levels, and reduced mortality.

## Results

### The effect of PF4 on plasma levels of NDPs in murine endotoxemia

We have previously observed that PF4-deficient mice (*Cxcl4*^−/−^) have increased mortality in LPS endotoxemia compared to animals that overexpress hPF4 (hPF4^+^)^29^. The infusion of wild-type (WT) mice with platelets obtained from hPF4^+^ animals improved survival^29^. We have recently found that PF4 binds to NETs, leading them to become physically compact and DNase resistant^27^. Therefore, we examined whether NET compaction with NDP sequestration contribute to the protective effect of PF4 in those studies. We measured NDP levels, including cfDNA, MPO, histone H3, and citrullinated histone H3 (cit-H3)) in *Cxcl4*^−/−^ and hPF4^+^ mice following LPS injection. *Cxcl4*^−/−^ mice had significantly higher levels of cfDNA and MPO than hPF4^+^ animals 3-8 hours post-LPS exposure (Figures 1A and 1B, respectively). H3 levels were on average higher in *Cxcl4*^−/−^ mice 8 hours following LPS injection, but this difference was not found to be significant. In comparison, levels of cit-H3, a more specific marker of NETosis^31^, were significantly elevated in *Cxcl4*^−/−^ animals (Figure 1C). We next explored whether infused hPF4 could recapitulate the protective effect of endogenous PF4 when administered via osmotic pumps or as a single intravenous (IV) bolus. We measured plasma levels of cfDNA and cfDNA-MPO complexes 6-hours post-LPS treatment and found both approaches led to decreased plasma NDPs (Figures 1D-1F, respectively).

**Figure 1.**
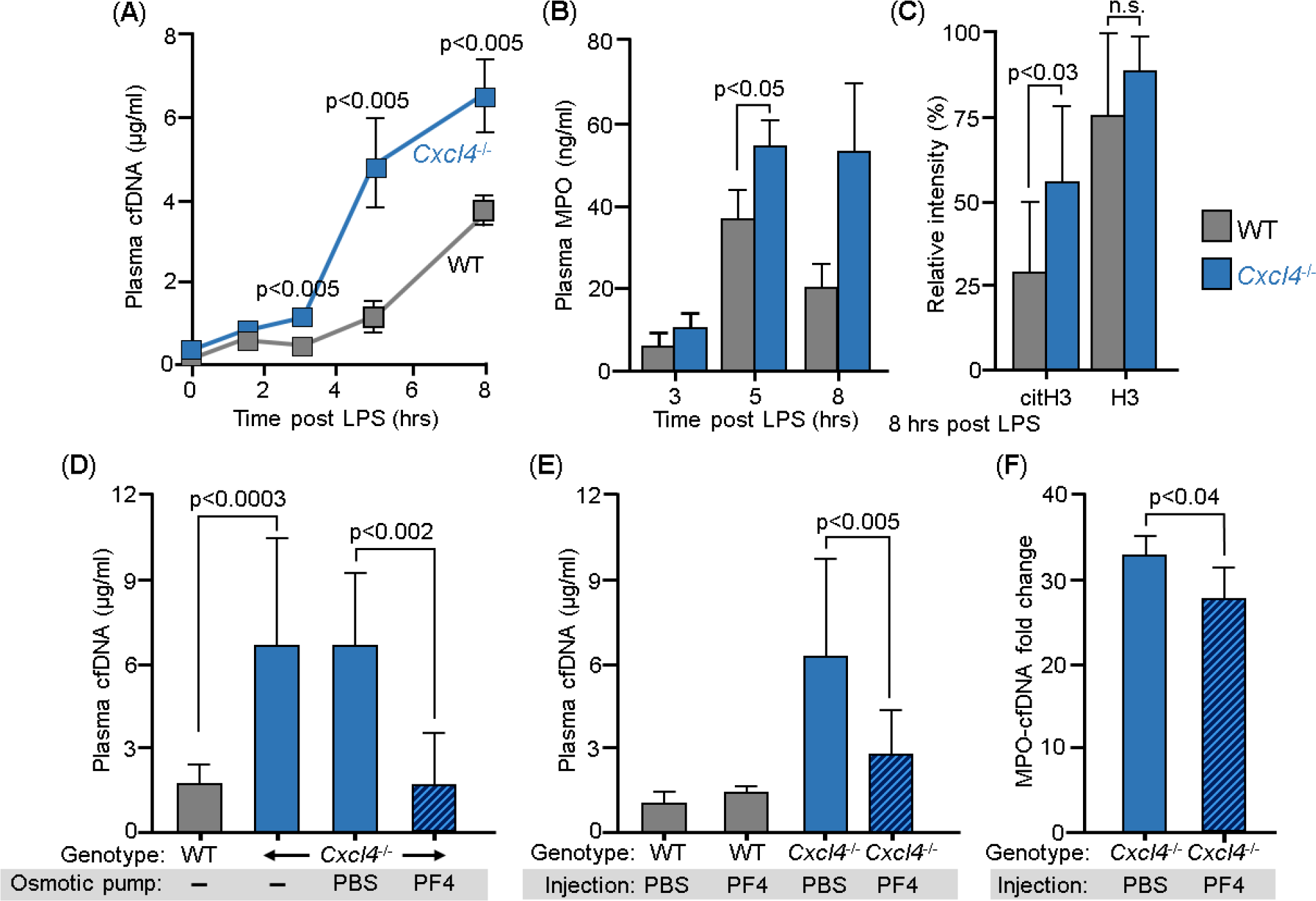
Effects of PF4 infusion on circulating NDP levels following LPS exposure in WT and *Cxcl4*^−/−^ mice. WT and *Cxcl4*^−/−^ mice received LPS (35 mg/kg, IP) and plasma samples were obtained at the indicated time points. (**A**) Mean plasma cfDNA levels are shown as ± 1 standard deviation (SD). N = 7-14 mice per arm. P values are indicated comparing WT and *Cxcl4*^−/−^ mice using a Mann-Whiney U test. (**B**) is the same as (A), but for MPO levels at 3 to 8 hours post-LPS. (**C**) Histone levels comparing western blot intensity to that of the positive control band. N = 9-10 mice per arm. P values are indicated comparing WT and *Cxcl4*^−/−^ mice by Mann-Whiney U test. (**D**) LPS studies comparing WT and *Cxcl4*^−/−^ mice following implantation of osmotic pumps containing PBS alone or 80 μg of hPF4. Mean ± 1 SD are shown N = 5-8 mice per arm. (**E**) is the same as (D), but following a single dose of hPF4 (80 μg, IV). N = 5-8 mice per arm. (**F**) LPS studies comparing plasma levels of cfDNA-MPO complex in *Cxcl4*^−/−^ mice that received tail vein injections containing PBS along or 80 μg of PF4. N=4-5 mice per arm. P values are indicated comparing WT and *Cxcl4*^−/−^ mice by Mann-Whiney U test.

### In vitro protective effects of PF4

In sepsis, end-organ injury involves damage to the microvasculature^32^. Therefore, we sought to determine if PF4-mediated compaction alters NETs’ ability to harm the endothelium. To that end, we stimulated isolated human neutrophils with LPS and resuspended the cells in media with and without hPF4 and flowed the samples through human endothelial umbilical vein cell (HUVECs)-lined microfluidic channels that had been stimulated with tumor necrosis factor (TNF)α. The neutrophils readily adhered to the endothelial monolayer, releasing NETs. The presence of PF4 led to NET compaction, although the cfDNA continued to fill the width of the channels (Supplement Figure 1). Flow was discontinued and the HUVECs were incubated with the NET containing media for 16-hours. Following the incubation, channels treated with PF4 contained a higher number of residual adherent endothelial cells, suggesting that PF4-bound NETs are less harmful to the endothelium (Figure 2A).

**Figure 2.**
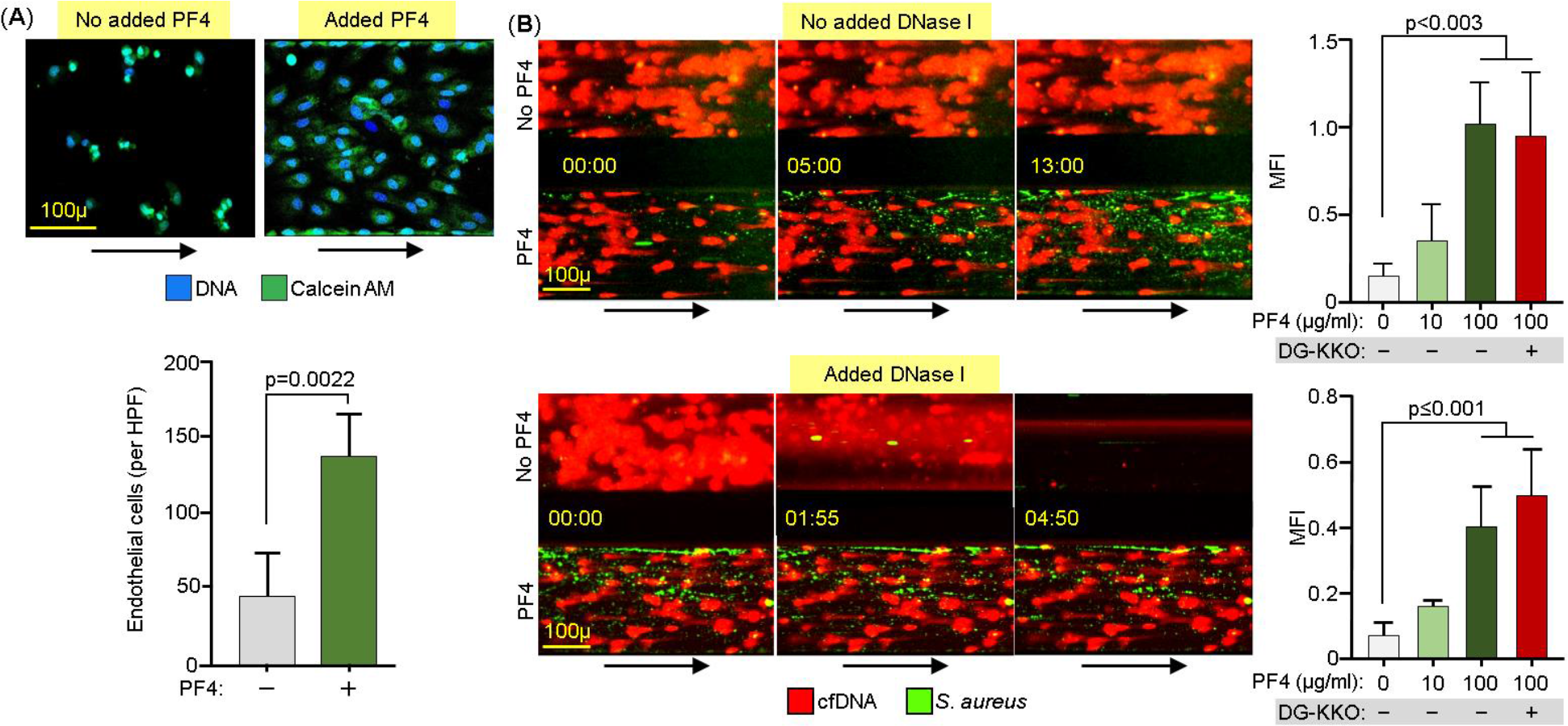
Effect of PF4 on endothelial cells and microbial entrapment by NETs in vitro. Channels lined with TNFα-stimulated HUVECs were infused with isolated human neutrophils treated with TNFα (1 ng/ml) with and without hPF4 (25 μg/ml). (**A**) Channels were incubated with the neutrophils for 16 hours after which the number of residual adherent endothelial cells was counted. At the top, representative images of remaining attached endothelial cells channels per condition. Size-bar and arrows indicating direction of flow are included. At the bottom, mean of endothelial cell counts in three 10x high-powered fields (HPFs) per condition ± 1 SD. Statistical analysis was performed using a Student’s test. (**B**) Left shows representative images of NET-lined channels infused with fluorescently-labeled *S. aureus* with observed bacterial capture. Size-bar and arrows indicating direction of flow are included. Right shows mean ± 1 SD quantified of the mean fluorescent intensity (MFI). For both left and right, top are studies without added DNase I and bottom in the presence of DNase I (100U/ml). N = 5-10 channels per condition. Analysis performed by a Kruskal-Wallis one-way ANOVA.

We then investigated how PF4 binding influences NETs’ ability to trap bacteria, another important way NETs impact outcome in sepsis^9^. We infused NET-lined channels with heat-inactivated, fluorescently-labeled *Staphylococcus (S) aureus* and used confocal microscopy to quantify bacterial capture. PF4-compacted NETs captured a significantly higher number of bacteria that adhered directly to the NET fibers (Figure 2B, top, Supplement Figure 2 and Supplement Video 1). Moreover, when these channels were infused with recombinant human DNase I, non-compacted, PF4-free NETs were rapidly digested leading to the liberation of captured bacteria (Figure 2B, bottom, Supplement Video 2). In contrast, during DNase infusion, compacted PF4-bound NETs remained intact and did not release their immobilized bacteria (Figure 2B, bottom, and Supplement Video 2).

### Studies of the in vitro effects of DG-KKO

By the time most patients present with the symptoms of sepsis, their platelets have been have already been stimulated to release large amounts of PF4^25^, limiting the therapeutic benefit of infusing additional PF4. Our prior observation that the HIT-like monoclonal antibody KKO enhances DNase resistance of hPF4-NET complexes^27^ raised the possibility that KKO may amplify the effect of endogenous PF4 and be of therapeutic benefit in sepsis. However, KKO fixes complement^33^, activates platelets^30^, and stimulates leukocytes^27,34^, inducing the development of a prothrombotic state in hPF4^+^/FcγRIIa-expressing mice^35^ and increasing mortality in murine endotoxemia (Figure 5D). We therefore modified KKO by deglycosylating the antibody^36^ as confirmed by liquid chromatography with tandem mass spectrometry (LC-MS/MS)^37^ (Supplement Figure 3). DG-KKO continued to bind hPF4-heparin and hPF4-NET complexes (Figures 3A and 3B, respectively), but had a reduced capacity to activate hPF4-exposed human platelets or induce thrombocytopenia in hPF4^+^/FcγRIIa^+^ mice (Figure 3C and 3D). In studies with NET-lined microfluidic channels, DG-KKO does not adhere to non-compacted NETs, but binds hPF4-compacted NETs (Figure 3E). Like KKO, DG-KKO also increased hPF4-NET resistance to DNase I under flow conditions in microfluidic channels (Figures 3F and 3G, Supplement Video 3) and did not impact the ability of PF4-NET complexes to capture bacteria (Figure 2B, Supplement Video 4).

**Figure 3.**
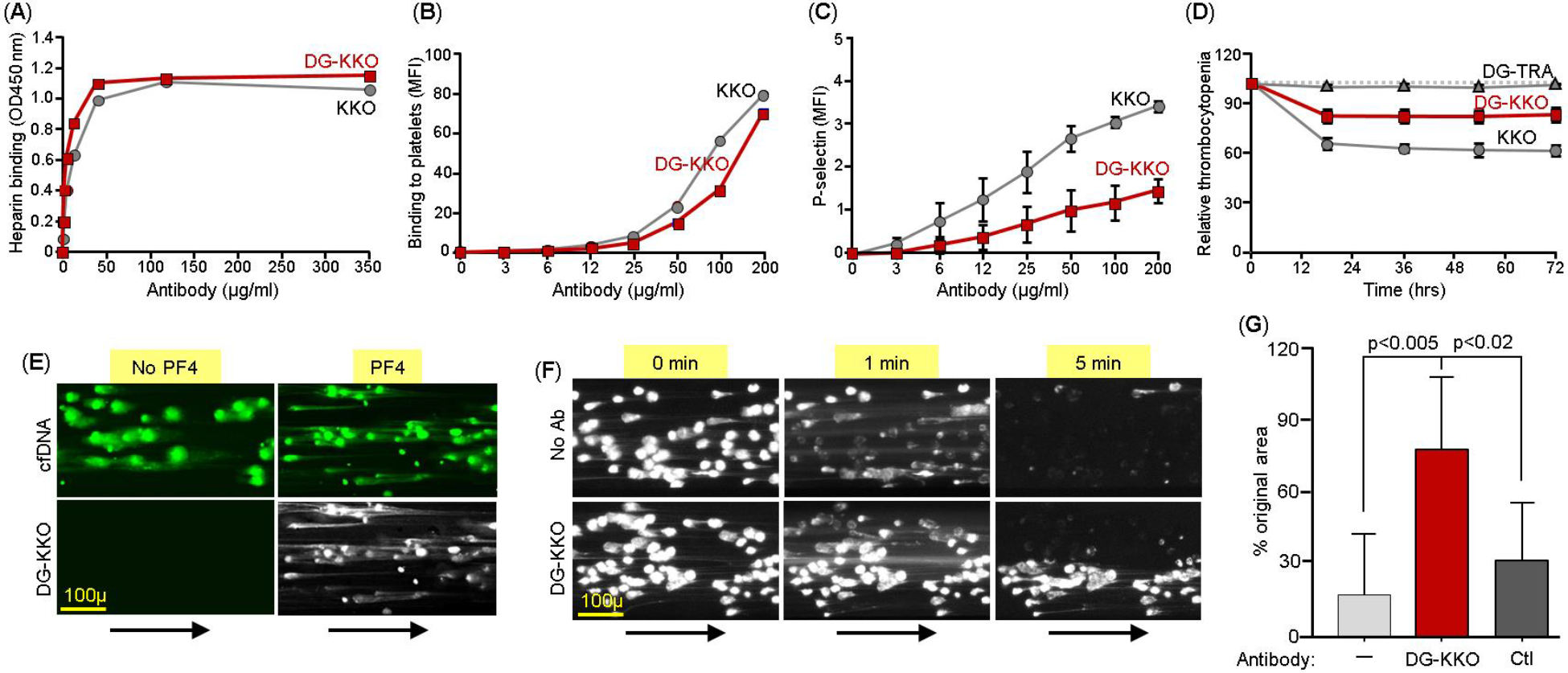
Binding of DG-KKO to PF4/NET complexes in vitro. (**A**) Graphs quantifying binding of increasing concentrations of KKO (gray) and DG-KKO (red) to heparin using fluorescent plate assay. (**B**) same as (A) but using flow cytometry to quantify antibody binding to the platelet surface. (**C**) Mean ± 1 SD of P-selectin MFI in human whole blood samples incubated with the indicated concentration of antibody, reflecting the degree of platelet activation. N=3. (**D**) Mean ± 1 SD of the % decrease in platelets counts in HIT mice injected with 400μg of the indicated antibody, measured every 12 hours for 3 days. N=3. (**E**) Representative confocal images of released NETs as in Figure 2 exposed to no PF4 or 6.5 μg/ml of PF4, labeled with the nucleic acid stain Sytox green (green) demonstrating change in morphology. The indicated channels were then infused with fluorophore-labeled DG-KKO (white). Size-bar and arrows indicating direction of flow are included. Image were obtained at 10x magnification. (**F**) Representative widefield images of adherent neutrophils as in (A), but in the presence of 100U/ml DNase I and 6.5 μg/ml PF4 ± 25 μg/ml of DG-KKO. (**G**) Mean ± 1 SD of the relative area of NETs compacted with PF4 (6.5 μg/ml) alone or PF4 plus either DG-KKP or a polyclonal anti-PF4 antibody control (Ctl) (each, 25 μg/ml KKO) post an infusion of DNase I (100U/ml, 3 minutes) compared to pre-infusion. N = 7-10 channels per condition. Comparative statistical analysis performed by Kruskall-Wallis one-way ANOVA.

### DG-KKO reduces plasma NDPs and improves outcomes in murine LPS endotoxemia

Based on these results, we investigated whether treatment with DG-KKO could improve outcomes in murine LPS endotoxemia. Thirty minutes post LPS-exposure, mice were given either DG-KKO, or a deglycosylated version of an isotype control antibody DG-TRA via tail vein injections. As both KKO and DG-KKO bind specifically to NET complexes containing hPF4 and not complexes with murine PF4 (Figure 3E and ^38^), hPF4^+^ mice were studied. We observed that LPS-exposed hPF4^+^ mice treated with DG-KKO were protected from the development of thrombocytopenia (Figure 4A). Moreover, animals treated with DG-KKO had lower levels of cfDNA and cfDNA-MPO complexes (Figures 4B and 4C), and the inflammatory cytokine monocyte chemoattractant (MCP)-1 (Figure 4D). DG-KKO treated animals did not have elevated plasma levels of thrombin anti-thrombin (TAT) complexes (Supplement Figure 4), indicating that DG-KKO stabilization of NETs did not lead to accelerated thrombin generation^39^. DG-KKO treated mice also had a more benign clinical course, demonstrated by significantly lower mean mouse sepsis scores (MSS^40^) at 12 hours (Figure 4E) and improved overall survival (Figure 4F). In parallel LPS studies of *Cxcl4*^−/−^ mice, DG-KKO did not exert a protective effect (Figures 4A to 4F), supporting that DG-KKO acts through an hPF4-dependent pathway in LPS endotoxemia.

**Figure 4.**
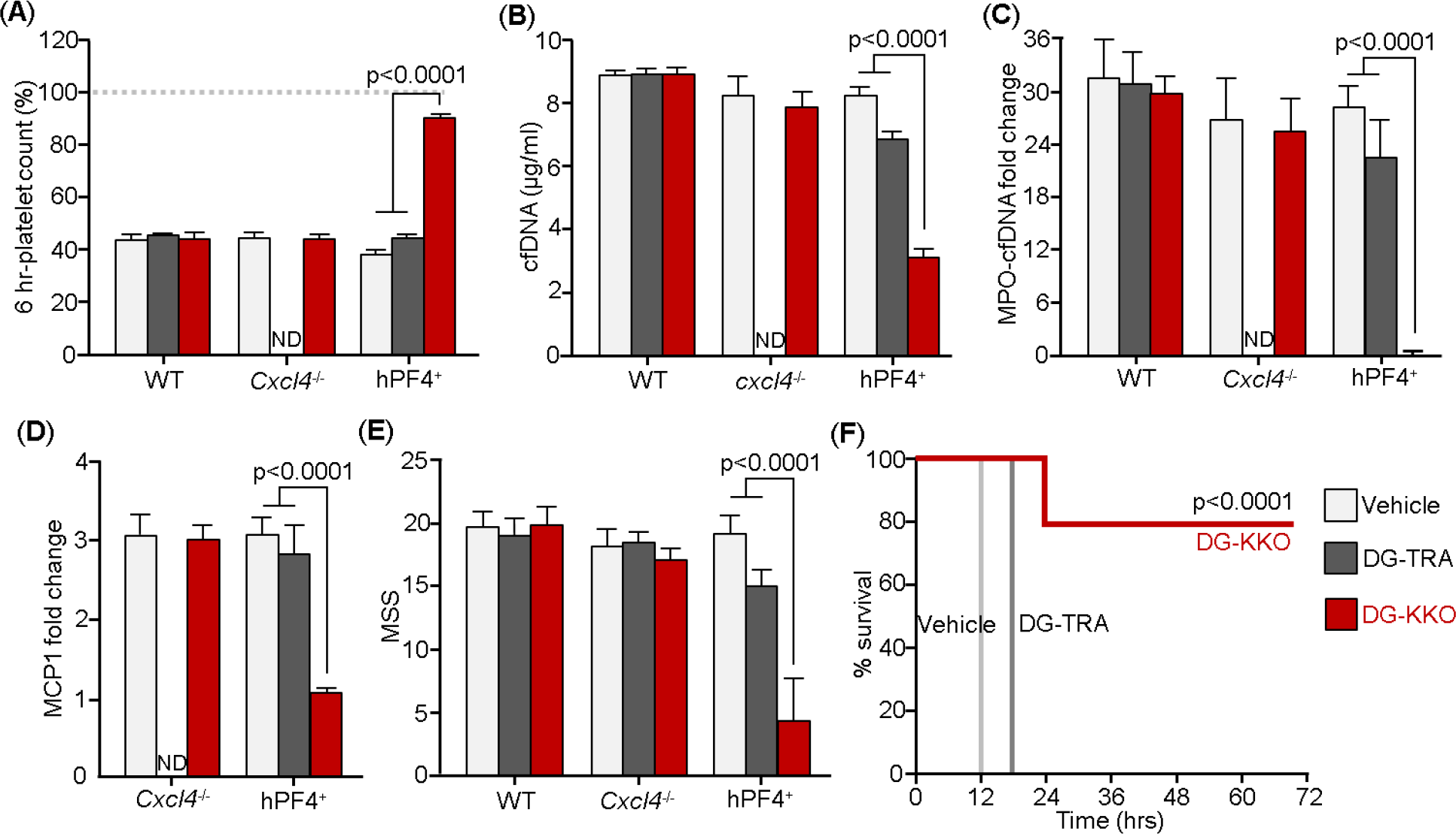
DG-KKO treatment in WT, *Cxcl4*^−/−^ and hPF4^+^ mice undergoing LPS endotoxemia. Mice were injection with LPS (35 mg/kg, IP). 30 minutes later they received tail vein injections containing vehicle alone or 5 mg/kg of DG-KKO or DG-TRA isotype control. 6 hours following LPS injection, a subset of mice were euthanized and IVC blood samples were collected. N = 3-10 in each arm. (**A**) Relative to baseline (dashed gray line), 6 hour-time point platelet counts. Mean ± 1 SD shown. ND = not done. Comparative statistical analysis was performed with Sidak’s multiple comparison test. (**B**) through (**E**) are the same as (**A**), but for cfDNA concentration, cfDNA-MPO fold change relative to negative control, MCP1 levels, and MSS, respectively. (**F**) are animal survival results for these LPS studies in the hPF4^+^ mice. Results were analyzed with Kaplan Meier survival analysis. See Supplement Figure 6 for survival results after DG-KKO in the WT and *Cxcl4*^−/−^ mice undergoing LPS endotoxemia.

To investigate the mechanism through which DG-KKO was acting, we repeated our murine LPS endotoxemia experiments with and without DNase I infusion, administered via tail vein injection 2 hours after LPS exposure. We observed that nuclease treatment led to decreased platelets counts in *Cxcl4*^−/−^ mice but had no impact on thrombocytopenia in hPF4+ animals (Supplement Figure 5A). *Cxcl4*^−/−^ mice treated with DNase had a significant increase in plasma cfDNA levels and higher mean sepsis scores (MSS)^41^ compared to untreated controls. The addition of DG-KKO did not exert a protective effect in these animals (Supplement Figure 5). In hPF4^+^ animals, treatment with DNase also led to a significant increase in cfDNA and cfDNA-MPO complexes. However, the NDP levels were significantly lower than those in *Cxcl4*^−/−^ animals, suggesting that the presence of hPF4 led to nuclease resistance (Supplement Figure 5B and 5C). hPF4^+^ mice treated with both DG-KKO and DNase had higher NDP levels and MSSs compared to animals treated with DG-KKO alone (compare Figure 5 and Supplement Figure 5), supporting our contention that NET-compaction rather than enhanced lysis improves outcome in LPS endotoxemia.

**Figure 5.**
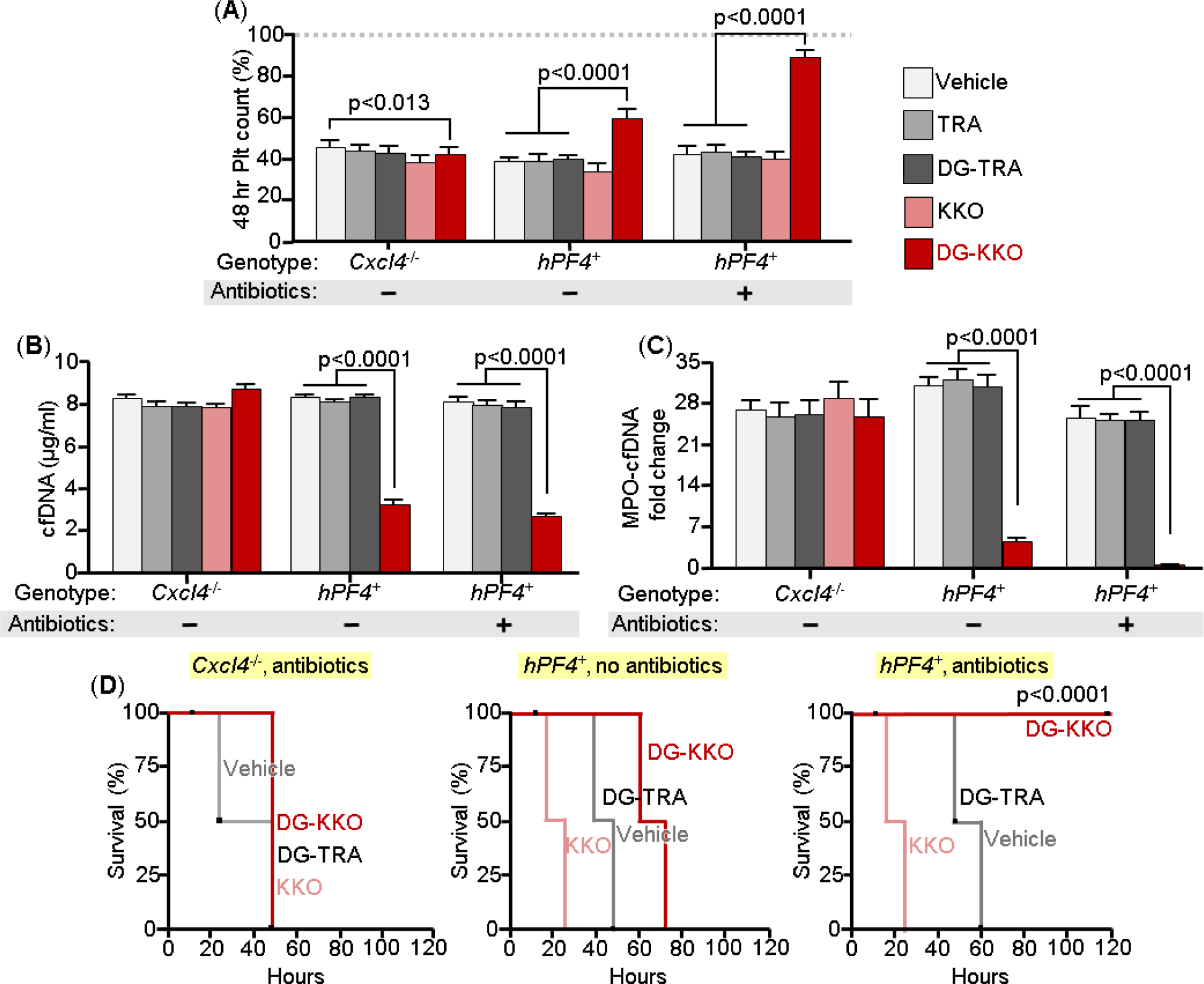
DG-KKO treatment in *Cxcl4*^−/−^ and hPF4^+^ mice undergoing CLP injury. All the mice underwent CLP procedure. Some mice, 1 hour following surgery, received an intradermal dose of the antibiotic ceftriaxone (100 mg/kg). Mice also were divided by therapeutic intervention, receiving either vehicle only or 5 mg/kg of KKO or DG-KKO or TRA or DG-TRA. After 48 hours, platelet counts and plasma NDP levels and survival were quantified. N = 10 animals per arm. Statistical analysis performed with Sidak’s multiple comparison t-test. (**A**) Relative platelet counts 6-hours post-CLP injury measured as in Figure 4A. (**B**) and (**C**) are as Figures 4B and 4C, respectively, but after a CLP injury. (**D**) are animal survival results for these CLP studies. Statistical analysis performed with Kaplan Meier survival analysis.

### DG-KKO improves outcomes in murine CLP

Although treatment with LPS recapitulates many of the aspects of sepsis, it relies on treatment with a bacterial toxin rather than infection with live pathogens. To determine if PF4 and DG-KKO mitigate NET-associated toxicities without impairing their positive effects in infection, we repeated our endotoxemia experiments using CLP to induce a polymicrobial sepsis model^42^. We performed these experiments in *Cxcl4*^−/−^ and hPF4^+^ mice with and without the administration of the antibiotic ceftriaxone given at a dose that does not improve outcome when used as a single agent to treat CLP^43^. We observed no protective effect of DG-KKO in *Cxcl4*^−/−^ mice following CLP (Figure 5). In contrast, in hPF4^+^ mice, DG-KKO prevented the development of thrombocytopenia after CLP (Figure 5A). These animals also had a reduction in plasma cfDNA and cfDNA-MPO complex levels (Figures 5B and 5C, respectively) and prolongation of survival (Figure 5D). Animals co-treated with ceftriaxone and DG-KKO survived CLP (Figure 5D), demonstrating that NDP sequestration may complement conventional antibiotic strategies in the treatment of polymicrobial sepsis.

### Co-administration of DG-KKO and hPF4 leads to improved outcomes in CLP in WT mice

We next investigated whether DG-KKO is protective when co-administered with exogenous hPF4 in WT mice. These studies were designed to further support our proposed model and also define the exogenous dose of hPF4 needed to provide clinical benefit in conjunction with an Fc-modified KKO. We observed that WT mice treated with hPF4 alone at doses of 20 mg/kg were not significantly protected following CLP. However, treatment with 40 mg/kg was associated with significantly less thrombocytopenia (Figure 6A), and lower plasma levels of cfDNA and cfDNA-MPO (Figures 6B and 6C, respectively). No protective effect was observed when animals were infused with DG-KKO and bovine serum albumin (BSA). However, mice treated with a combination of DG-KKO and the lower dose of hPF4 (20 mg/kg) did well with the highest platelet counts (Figure 6A), the lowest plasma NDP levels (Figures 6B and 6C), and improved MSSs (Figure 6D). These results indicate it may be feasible to replicate our murine studies in larger animal models wherein their PF4-NET complexes are not recognized by KKO, using a combination of exogenous hPF4 and DG-KKO to achieve maximal effect. These results also suggest that co-infusion of hPF4 with DG-KKO may be a superior therapeutic option to either treatment alone.

**Figure 6.**
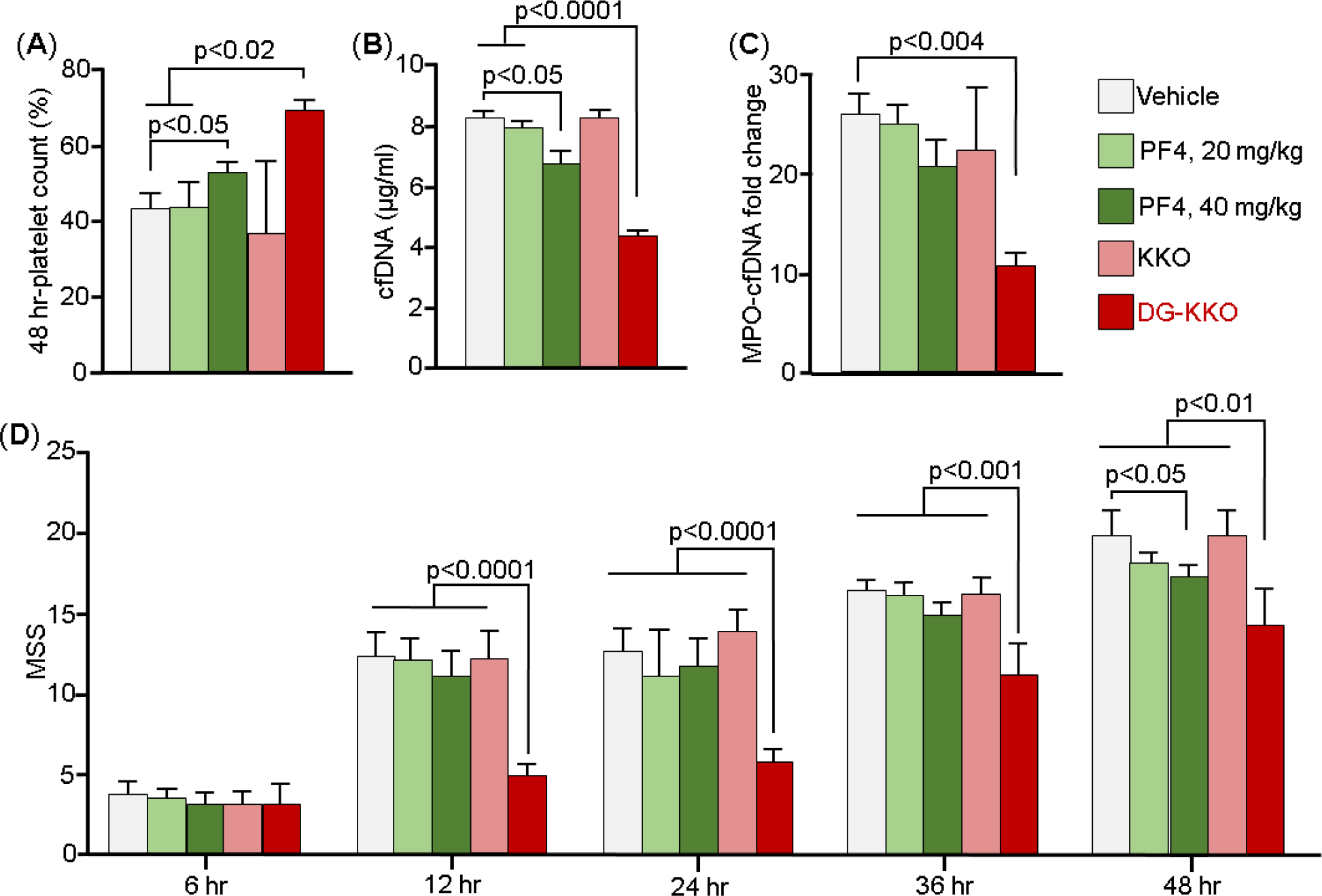
Treatment with PF4 or PF4 plus KKO or DG-KKO in WT mice undergoing CLP injury. CLP injuries were done as in Figure 5, but with WT mice that either received vehicle, PF4 at either 20 or 40 mg/kg, or combination therapy of PF4 (20 mg/kg) plus either 5 mg/kg KKO or DG-KKO. N = 10 animals per arm. (**A**) Relative platelet counts as in Figure 5A. (**B**) and (**C**) are as in Figures 5B and 5C, respectively. (**D**) shows MSS in these studies for up to 48 hrs after the injury. Mean ± 1 SD shown. In (A) through (D), statistical analysis was performed with Sidak’s multiple comparison t-test.

## Discussion

In studies designed to investigate the role of neutrophils in the prothrombotic nature of HIT, we observed that PF4, a chemokine known to cross-aggregate polyanions like heparin^26^, also binds to NETs, presumably via the polyanionic backbone of their DNA, causing them to become physically compact. It has recently been shown that PF4-cfDNA complexes form in vivo during systemic sclerosis^28^ and it is likely that they also occur during conditions such as sepsis that precipitate both NETosis and platelet degranulation^22^. We have recently shown that PF4 binding enhances NET resistance to lysis by DNase I and microbial nucleases without displacing histones^27^. We now show that PF4 protects the endothelium from damage induced by NETs likely by limiting the release of histone as well as other toxic NDPs. We also demonstrate that PF4, which has previously been shown to bind to the surface of bacteria via electrostatic interactions^44^, enhances the ability of NETs to entrap *S. aureus*.

Cumulatively, our in vitro studies demonstrate that PF4 binding decreases NET toxicity by inhibiting NDP release and enhances bacterial entrapment by improving microbial capture and limiting susceptibility to DNase lysis (Figure 7). Based on these results, we speculated that PF4-mediated NET compaction may be a conserved function that increases NET functionality while decreasing collateral host tissue damage in conditions such as sepsis in which overwhelming NET release occurs ^45^. This theory in part stemmed from the observation that many species of bacteria release nucleases as a virulence factor to evade capture by NETs^46,47^. It was also based on a prior study conducted by our group in which we observed that PF4 expression led to improved survival in murine LPS endotoxemia^29^.

**Figure 7.**
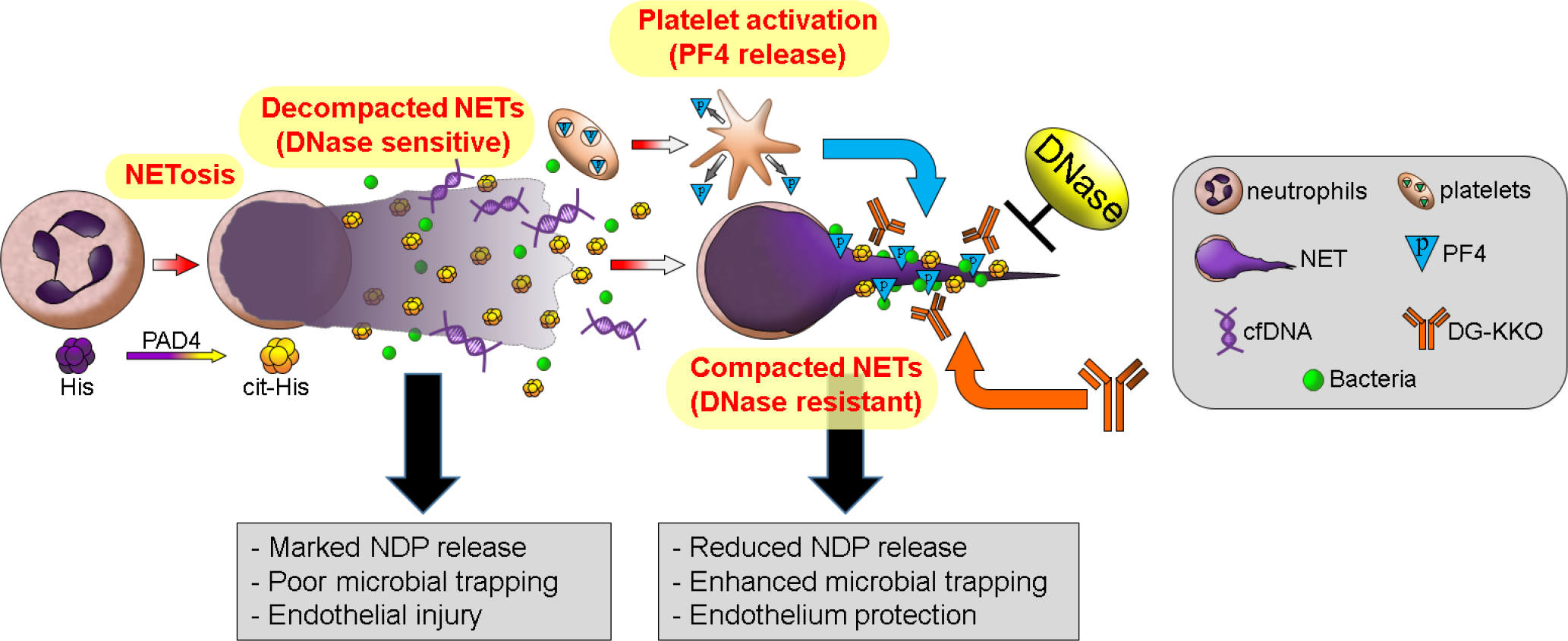
Schematic of PF4- and KKO-mediated NET compaction and NDP sequestration. Neutrophils release NETs after the enzyme peptidyl arginine deiminase 4 (PAD4) citrullinates histones (His) to become cit-His. Toxic NET degradation products (NDPs) such as cfDNA and histones, are released from NETs after DNase I digestion, leading to platelet activation and endothelial damage. At the same time, activated platelets release PF4 that binds to NETs, causing them to become dense, compact and partially resistant to DNase I. It also enhances their ability to capture bacteria. DG-KKO binds to PF4-NET complexes, enhancing their resistance to DNase I, sequestrating NDPs, and improving outcome.

We considered whether treatment with exogenous PF4 might improve outcomes in sepsis. However, by the time they become clinically ill, most septic patients are likely to have a high level of platelet activation and PF4 release^48,49^. As platelet counts and PF4 levels per platelet vary greatly in the general population^50,51^, we were concerned that supplemental PF4 infusions may vary widely in therapeutic efficacy.

An alternative treatment strategy would be an intervention that stabilizes interactions between NETs and endogenous PF4. It has previously been shown that PF4 binding to DNA aptamers exposes HIT antigenic sites^52^. Therefore, it was not surprising that the HIT-like monoclonal antibody KKO bound specifically to PF4-NET complexes, further enhancing their DNase resistance^27^. We now show that KKO binding did not interfere with the ability of PF4-NETs to entrap *S aureus*. KKO, or another HIT-like antibody, might enhance PF4 efficacy by further stabilizing PF4-NET complexes, decreasing NDP release and protecting the endothelial lining, while maintaining bacterial entrapment by NETs (Figure 7). However, KKO, is an IgG2bκ antibody that activates Fc receptors and induces complement-dependent cytotoxicity (CDC)^53^. Therefore, unmodified KKO itself was likely to be injurious in sepsis. In fact, in CLP studies of hPF4^+^ mice, animals treated with unmodified KKO showed accelerated demise (Figure 5D). IgG2bκ antibodies are also resistant to cleavage to form F(ab)2 fragments^54^, so we instead treated KKO with EndoS to generate degylcosylated DG-KKO. Deglycosylated antibodies have been used in the treatment of immune-mediated disease. Infusion with IdeS, a bacterial protease that cleaves the hinge region of heavy chain IgG and abolishes its ability to bind to FcγR, reduces the pathogenicity of autoantibodies in HIT, immune thrombocytopenia, and solid organ transplant^55–57^. EndoS has also been used to deglycosylate pathogenic anti-aquaporin-4 antibodies in neuromyelitis optica, converting them to therapeutic blocking antibodies^58^.

In vitro studies confirmed that DG-KKO retained the ability to bind to PF4-NET complexes, increasing their resistance to DNase without interfering with their ability to capture bacteria. In vivo results demonstrate that DG-KKO amplifies the protective effect of PF4 in sepsis, leading to a reduction in thrombocytopenia, decreased plasma levels of NDPs, and improved survival in both murine LPS endotoxemia and CLP sepsis models. These benefits were only observed in hPF4^+^ animals, confirming that DG-KKO acts through an hPF4-dependent mechanism.

Two alternative NET-based approaches have been proposed to improve outcome in sepsis. One is to infuse DNases into septic animals. Our experiments with DNase I infusions are consistent with the results of prior murine CLP studies in which the early administration of DNase leads to increased inflammation and decreased survival^24^. We found that there was a greater increase in plasma NDP levels following DNase infusion in *Cxcl4*^−/−^ mice compared to hPF4^+^ mice, supporting that PF4 protects NETs from DNase lysis in vivo. Moreover, DG-KKO continues to be protective in hPF4^+^ mice even following DNase infusion.

The other NET-based therapeutic strategy that has been studied is the blockade of NETosis through peptidyl arginine deiminase (PAD) 4 inhibition. While PAD4 blockade may prove to be a useful treatment for autoimmune disease, it is an imperfect strategy in the treatment of sepsis as NETs entrap bacteria and may enhance their killing^59^. Therefore, preventing NETosis could lead to increased bacterial dissemination. To date, the results of studies investigating PAD4 inhibition in murine models of sepsis have been mixed with some studies showing minimal negative effects^21^ and others revealing increased vulnerability to infection^9,60^. The second problem with blocking NETosis is that by the time a patient presents with sepsis, their neutrophils have likely already released a large burden of NETs. In clinical studies of patients with septic-shock, levels of cfDNA-MPO were found to be highest at the time of admission^19^. Therefore, even prompt administration of a PAD4 inhibitor is likely to be too little too late in the treatment of most patients with sepsis.

Our results suggest that in sepsis, NET compaction and NDP sequestration may be superior to PAD4 blockade or DNase infusion. We have found that it is protective in both sterile LPS endotoxemia and CLP polymicrobial sepsis. Whether it may also be effective in other conditions characterized by NET release including disseminated intravascular coagulation^61^, sickle cell disease^62^, antiphospholipid antibody syndrome^63^, rhabdomyolysis^64^ and burns injuries^65^ is unclear, and would need to be tested in comparison to alternative strategies.

In summary, we believe that NET compaction and NDP sequestration by infusions of PF4 and/or an Fc-modified HIT antibody provide new insights into the mechanism by which NETs contribute to the pathophysiology of sepsis. The continued study of this pathway may lead to the development of targeted therapeutic strategies to improve outcome in sepsis and perhaps other disorders associated with excess NET release.

## Methods

### Antibodies studied and recombinant hPF4

KKO is a mouse IgG 2bκ anti-hPF4/heparin monoclonal antibody^30^. TRA is a monoclonal IgG isotype control antibody^30^. Both KKO and TRA were purified from hybridoma supernatants^30^. KKO and TRA were deglycosylated using EndoS (IgGZERO, Genovis)^66^ adding 1 unit IgGZERO to 1 μg of KKO or TRA incubated at 37°C for 2 hours^67^. Liquid Chromatography with tandem mass spectrometry (LC-MS/MS)^37^ was used to confirm removal of the Fc glycan moieties from KKO (Supplement Figure 3).

Recombinant hPF4 was expressed in S2 cells^68^, and purified using affinity chromatography and protein liquid chromatography as previously described^68^. The end-product purity was found to be endotoxin free and was tested for size distribution by gel electrophoresis (SDS-PAGE). Its ability to bind to heparin and yield an immunogenic product was confirmed by ELISA as previously described^68^.

### NET-lined microfluidic channel studies

NET-lined channels were generated as described^27^. Human neutrophils were isolated as in the Supplement Method. These cells (2 × 10^6^ cells/ml) were incubated with 1 ng/ml TNFα (Gibco) and allowed to adhere to channels coated with fibronectin (Sigma-Aldrich)^35^. All infusions in these studies were done at 2-5 dynes/cm^2^. After the neutrophils were adherent, channels were incubated with 100 ng/ml phorbol myristyl acetate (PMA, Sigma-Aldrich) in Hank’s buffered salt solution (HBSS with calcium and magnesium, Gibco) overnight at 37°C. Release of cfDNA was visualized with 1 μM SYTOX green or orange (Thermo Fisher Scientific) also suspended in HBSS. Channels were then infused with hPF4 (0-100 μg/ml). Some channels containing PF4-NET complexes were then incubated with 25 μg/ml of KKO, DG-KKO, or polyclonal anti-PF4 antibody (25μg/ml, Abcam) for 1 hour at 37°C. NET digestion studies were carried out by infusing the channels with 100 U/ml DNase I, Sigma-Aldrich) as described^27^. KKO and DG-KKO were labeled with Alexa Fluor 647 (Thermo Fisher Scientific) prior to NET channel infusion. NET complexes were imaged with a Zeiss LSM 710 laser-scanning confocal microscope. Experiments were performed using a BioFlux 200 Controller (Fluxion)^69^. The BioFlux channels were visualized with an Axio Observer Z1 inverted Zeiss microscope equipped with a motorized stage and an HXP-120 C metal halide illumination source. The microscope and image acquisition were controlled by BioFlux Montage software with a MetaMorph-based platform (Molecular Devices). Data were analyzed using ImageJ open source image processing software^27^.

In some NET-lined channel studies, Alexa Fluor 488-labeled, heat-killed *S aureus* (40 μg/ml, Molecular Probes) suspended in HBSS were infused for 30 minutes at 2 dynes/cm^2^. These NET channels had previously been treated with HBSS alone ± hPF4 at concentrations of 10 or 100 μg/ml. Bacterial adhesion was quantified over time as measured by an increase in channel mean fluorescent intensity. Some channels were then infused with DNase I (100U/ml, Sigma-Aldrich) for 15 minutes. The channels were then washed and infused with Cellfix (BD Biosciences) for 15 minutes prior to imaging with a Zeiss LSM 710 laser-scanning confocal microscope.

### Endothelialized channel microfluidic studies

Similar studies were done as described in the above NET-lined channel studies, but the channels were first lined with human umbilical vein endothelial cells (HUVECs, Lonza) at passage 3-4 (5 × 10^6^ cells per channel), seeded onto fibronectin-coated channels, and then cultured at 37°C under 5% CO_2_ in endothelial cell growth media (Lonza) for 2-3 days to become confluent 35. HUVECs were incubated with TNFα (10 ng/ml, Thermo Fisher Scientific) to simulate inflammation. Cell nuclei were labelled with Hoescht 33342 (Thermo Fisher Scientific). Isolated neutrophils were fluorescently labeled with 2-mM calcein AM (Thermo Fisher Scientific) for 15 minutes at 37°C ^70^. The samples were then recalcified with CaCl_2_ (11 mM, final concentration), stimulated with LPS (1 ng/ml, Sigma-Aldrich), suspended in endothelial cell growth media containing with SYTOX orange (1 μM, Thermo Fisher Scientific) to visualize DNA and flowed through the HUVEC-lined channels at 5 dynes/cm^2^ for 30 minutes, during which time the neutrophils adhered to the HUVEC monolayer and released NETs. After completion of flow, the channels were incubated with the NET-containing buffer at 37°C under 5% CO_2_ overnight. Channels were then infused with Cellfix for 15 minutes. Image analysis was performed as described for the NET-lined channel studies.

### Mice studied

Mice were either WT C57Bl6 or littermate mice deficient in murine PF4 (*Cxcl4*^*−/−*^) or transgenic for platelet-specific expressed hPF4 on a *Cxcl4*^*−/−*^ background (termed hPF4^+^ mice)^71^ as murine PF4 is not recognized by KKO and by most patient HIT antibodies^38^. hPF4^+^ mice that also transgenically expressed human FcγRIIA in a tissue-specific fashion^72^ were used in a HIT passive immunization model^68^. Genetic modifications in all these mice were confirmed by PCR analyses^73^. All mice were studied between 10-20 weeks of age. There was an equal distribution of gender in the mice generated, and animals of each gender were randomly studied in the LPS and CLP sepsis, and the HIT passive immunization murine model.

### LPS endotoxemia sepsis model

Baseline platelet counts from retro-orbital blood samples were measured using the Hemovet 950 (Drew Scientific). On the following day, WT, *Cxcl4*^−/−^ or hPF4^+^ mice received an intraperitoneal (IP) injection of LPS at a dose of 35 mg/kg. 30 minutes later, the mice received a tail vein injection containing of either vehicle alone or KKO or DG-KKO or TRA or DG-TRA at doses of 5 mg/kg diluted in 200 μl of phosphate buffered saline (PBS with CaCl_2_ and MgCl_2_, Gibco). Five hours after LPS treatment, blood was collected from the retro-orbital plexus and platelet counts were measured. A subset of mice was treated with IP with recombinant DNase I (20 mg/kg, Sigma-Aldrich) 2 hours after LPS injection^24^. Six hours following LPS injection, a subgroup of animals was sacrificed, blood was collected from the inferior vena cava (IVC), and plasma was isolated. cfDNA was measured using a SYTOX fluorescent plate assay^74^, cfDNA-MPO was measured using a previously described ELISA^75^, and MCP-1 levels were quantified with western blot, using anti-MCP-1 (Cell Signaling Technology, #2027). Another subgroup had MSS calculated and survival noted every 12 hours as described^74^ for up to 72 hours. When MSS exceeded 18, the animals were euthanized in accordance with IACUC protocol.

### CLP polymicrobial sepsis model

CLP-induced sepsis was induced using a modified version of a published method^76^ in WT, *Cxcl4*^−/−^ and hPF4^+^ mice. Mice were anesthetized using IP ketamine (150 mg/kg) and xylazine (10 mg/kg) prior to abdominal surgery to expose the cecum^77^. The cecum was exteriorized and ligated with a 6.0 silk suture (6-0 PROLENE, 8680G; Ethicon) placed below the ileo-cecal valve and perforated with a 21-gauge needle (BD Biosciences) to induce mid-grade lethal sepsis^78–80^. After removing the needle, a small amount of feces was extruded. The cecum was relocated, and the fascia, abdominal musculature, and peritoneum were closed via simple running suture. The control mice were anesthetized and underwent laparotomy without puncture or cecal ligation. Following the procedure, 1 ml of saline was administered subcutaneously for fluid resuscitation^81^, and the animals received a tail vein injection of PBS, KKO, DG-KKO, TRA or DG-TRA at doses of 5 mg/kg. In some WT mice studies, the animals were given either 20 or 40 mg/kg of hPF4 by tail vein injection. All animals were treated with 0.05 mg/kg buprenorphine every 12 hours to maintain analgesia. In some experiments, ceftriaxone was injected intradermally following surgery (100 mg/kg, Tocris Bioscience). Six hours after surgery, in a subset of mice, blood was collected from the retro-orbital plexus and platelet counts were measured with a HemaVet 950FS (Drew Scientific). Another subset of animals was sacrificed at 48 hours, and blood was obtained from the IVC to measure NDP levels. Another subgroup had MSS calculated and survival noted every 12 hours as described^74^ for up to 120 hours.

### Statistical analysis

Normal distribution was tested using the D’Agostino & Pearson normality test. When the data was not found to be normally distributed, differences between 2 groups were compared using a Mann-Whitney *U* test. Differences between more than 2 groups were determined with a Kruskall-Wallis test or with Sidak’s multiple comparisons tests as appropriate. Multiplicity corrected P values are reported for multiple comparisons. Statistical analyses were performed using Microsoft Excel 2011 and GraphPad Prism 7.0 (GraphPad Software). Differences were considered statistically significant when *P* values were ≤0.05.

### Study approval and additional methodology

Animal procedures were approved by the Institutional Animal Care and Use Committee (IACUC) at the Children’s Hospital of Philadelphia (CHOP) in accordance with NIH guidelines and the Animal Welfare Act. Anonymized human blood was collected after signed, informed consent was provided by healthy donors. Approval for the use of human blood was obtained from CHOP’s Human Review Board in accordance with Declaration of Helsinki Principles.

### Additional methodologies

Supplement Methods has a description of antibodies and other labeled probes, isolation of human neutrophils, and biosassaysbioassays for NDPs.

### Authorship

KG and AS carried out and evaluated these studies. KG and AS prepared the first draft and subsequent revisions of this manuscript. LR and MAK provided research guidance. SS performed the mass spectroscopy studies. MP provided overall project organization and direction, data interpretation, and manuscript preparation. None of the authors have any financial or related disclosures to make with respect to this manuscript.

## Acknowledgments

This work was supported by the National Heart Lung and Blood Institute grants (R01 HL139448 (MP) and T32HL007150 (KG)) and by an American Society of Hematology Fellow Research Award (KG).

