## Supplement for "Neutrophil extracellular trap stabilization leads to improved outcomes in murine models of sepsis"

**Supplement Methods**

*Description of antibodies and other labeled probes*

KKO, an anti-PF4/heparin monoclonal antibody, and TRA were purified from hybridoma supernatants. All antibodies were either labeled using Alexa Fluor antibody-labeling kits according to manufacturer's instructions or species-appropriate Alexa Fluor-conjugated secondary antibodies (all from Thermo Fisher Scientific). Monocyte chemoattractant protein (MCP)-1 was quantified using western blot with rabbit anti-MCP-1 (Cell Signaling Technology, cat no. 2029S) and mouse anti-GAPDH (Cell Signaling Technology, cat no. 97166T) as a control. Mouse TAT complexes were quantified using a commercially available ELISA (Assaypro).

*Isolated of human neutrophils*

Human blood was collected after informed consent from healthy aspirin-free volunteers through a 19-gauge butterfly needle into 4 ml vacutainers coated with 7.2 mg of ethylenediaminetetraacetic acid (Becton Dickinson). Blood samples were kept at room temperature and used within 1 hour. Neutrophils were isolated with negative bead selection using the MACSxpress Neutrophil Isolation Kit (Miltenyi Biotec). The supernatant was collected and centrifuged at 300g for 10 minutes, and the cell pellet was resuspended in 2 ml ammonium-chloride-potassium (ACK) lysing buffer (Thermo Fisher Scientific) for 5 minutes to lyse erythrocytes, after which the cells were washed with 10 mls of Hank's balanced salt solution (HBSS, Thermo Fisher Scientific), and then resuspended in HBSS containing calcium chloride (1.3 mM) and then manually counted, with trypan blue dye exclusion (Gibco) for cell viability.

*Bioassays for NDPs*

Released cfDNA was visualized using SYTOX green or orange nucleic acid stain (Thermo Fisher Scientific) and quantified using a previously described Sytox green fluorescent plate assay[1]. Briefly, plasma samples were diluted 1:10 in HBSS samples were incubated with SYTOX green (1 µM) and 50 µl were plated in duplicate in a solid black opaque 96-well plate (Corning). A baseline reading was obtained with fluorescence spectrometry, excitation 485nm and emission 537nm (SpectraMax M2, Molecular Devices). Rabbit anti-cit-H3 (Ab5103, Abcam) was used in western blot to quantify cit-H3[2]. Myeloperoxidase was quantified using a commercial ELISA kit cfDNA (Biolegend). MPO-DNA complex levels measured as previously described[1].

### Supplement Videos

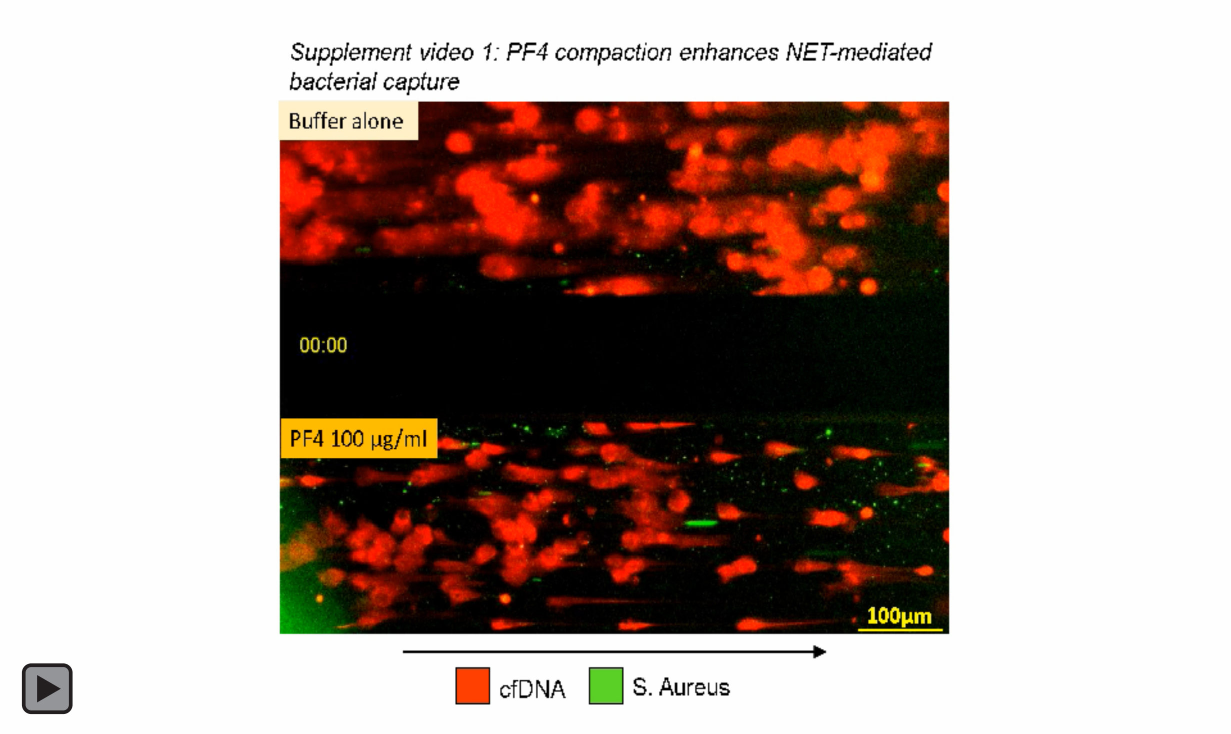


**Supplement Video 1. PF4 binding enhances NET-mediated capture of *S aureus* bioparticles in a microfluidic system.**

NETs adherent to fibronectin-coated microfluidic channels were labeled with SYTOX orange and infused with HBSS alone (top channel) or HBSS containing PF4 100 μg/ml to induce compaction (bottom channel). A video was obtained when these channels were infused with Alexa Fluor 488-labeled, heat-killed *S aureus* (40 μg/ml) suspended in HBSS for 30 minutes at 2 dynes/cm$^2$. An arrow indicating the direction of flow is included.

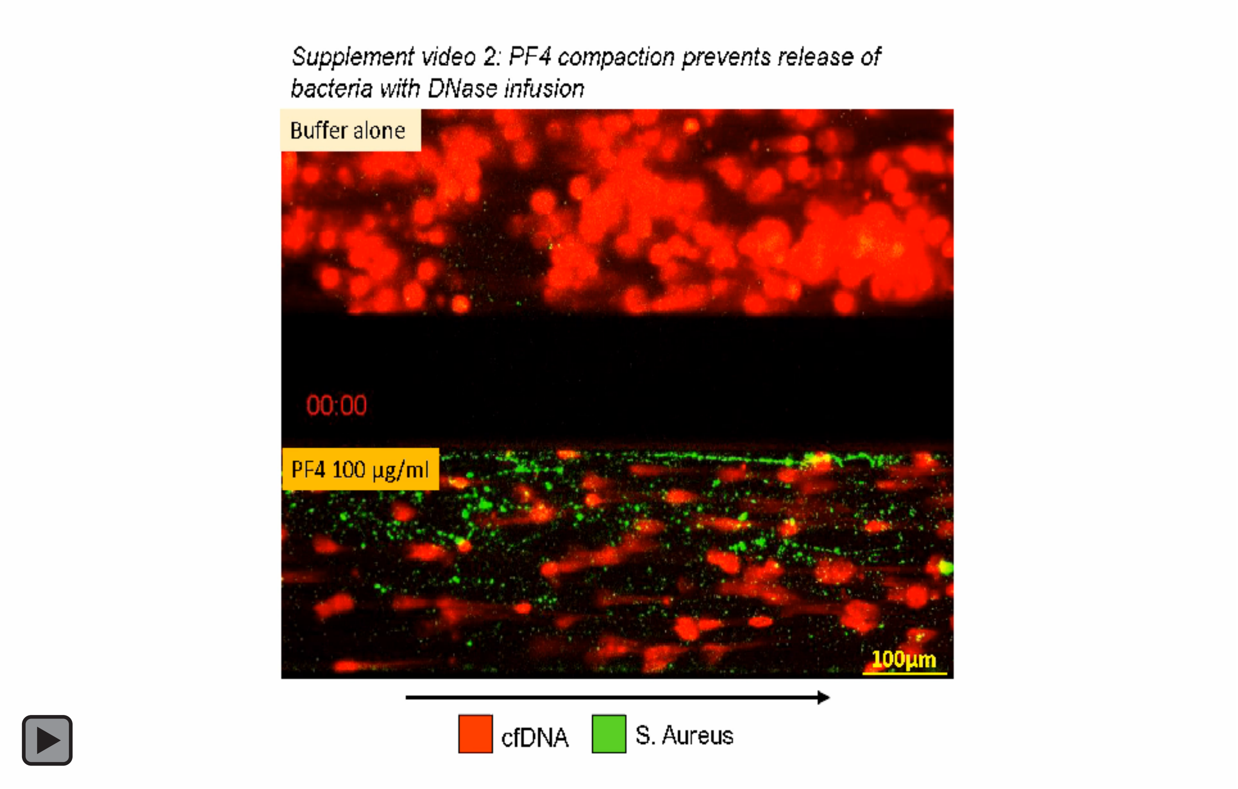


**Supplement Video 2. PF4 binding prevents the release of S aureus bioparticles from compacted NETs during DNase I infusion.**

NET-lined channels ± PF4 100 µg/ml, previously infused with *S aureus* bioparticle for 30 minutes, are now infused with DNase I 100 U/ml at the same flow rate (2 dynes/cm$^2$) over 3 minutes. An arrow indicating the direction of flow is included.

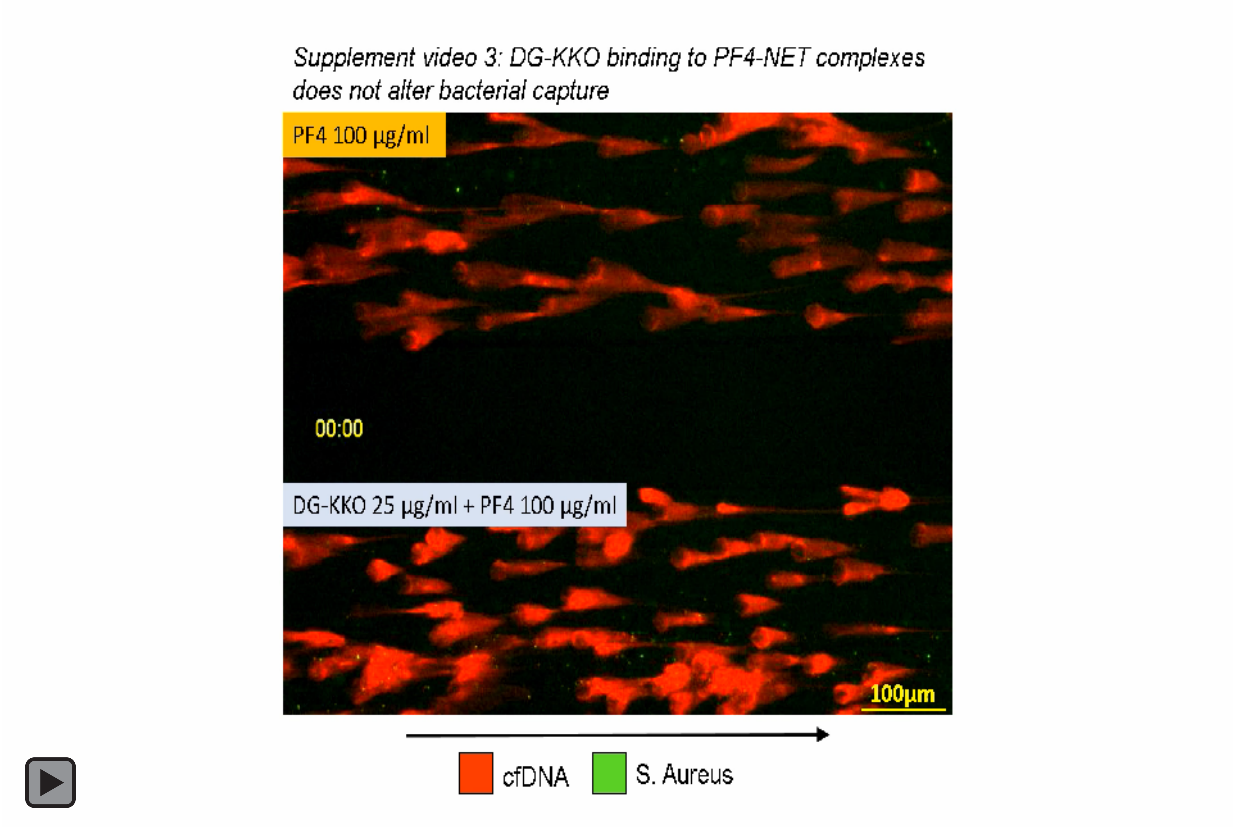


**Supplement Video 3. DG-KKO binding enhances PF4-NET complexes resistance to digestion with DNase 100U/ml**

NETs (stained light grey for released chromatin) were incubated with an intermediate concentration of PF4 (6.5 µg/ml). These PF4-NET complexes were exposed to buffer alone (top channel) or DG-KKO at 25 µg/ml (bottom channel) and then simultaneously infused with DNase I (100U/ml). An arrow indicating the direction of flow is included.

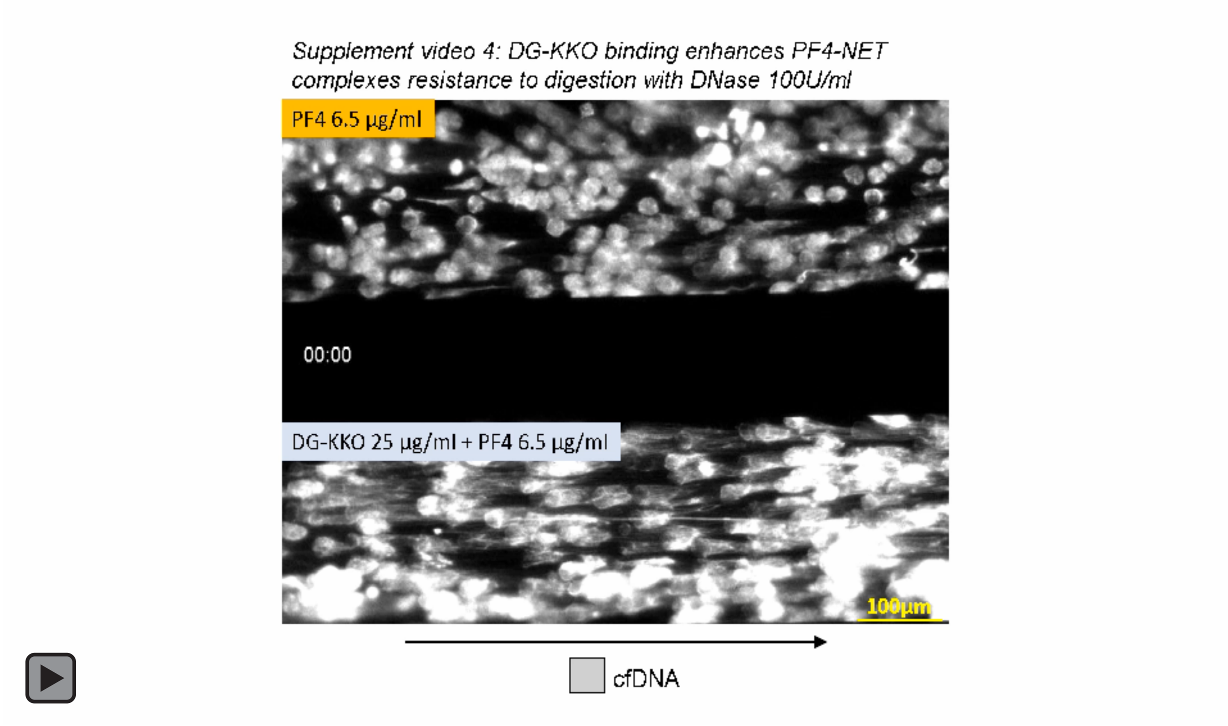


**Supplement Video 4. DG-KKO binding to PF4-NET complexes does not decreased bacterial capture**

NET-lined channels were infused with PF4 100 µg/ml for 30 minutes. The channels were subsequently incubated with HBSS alone (top channel) or KKO at 5 µg/ml. Channels were then infused with *S aureus* bioparticle for 30 minutes at 2 dynes/cm$^2$. An arrow indicating direction of flow is included.

**Supplement Figures**

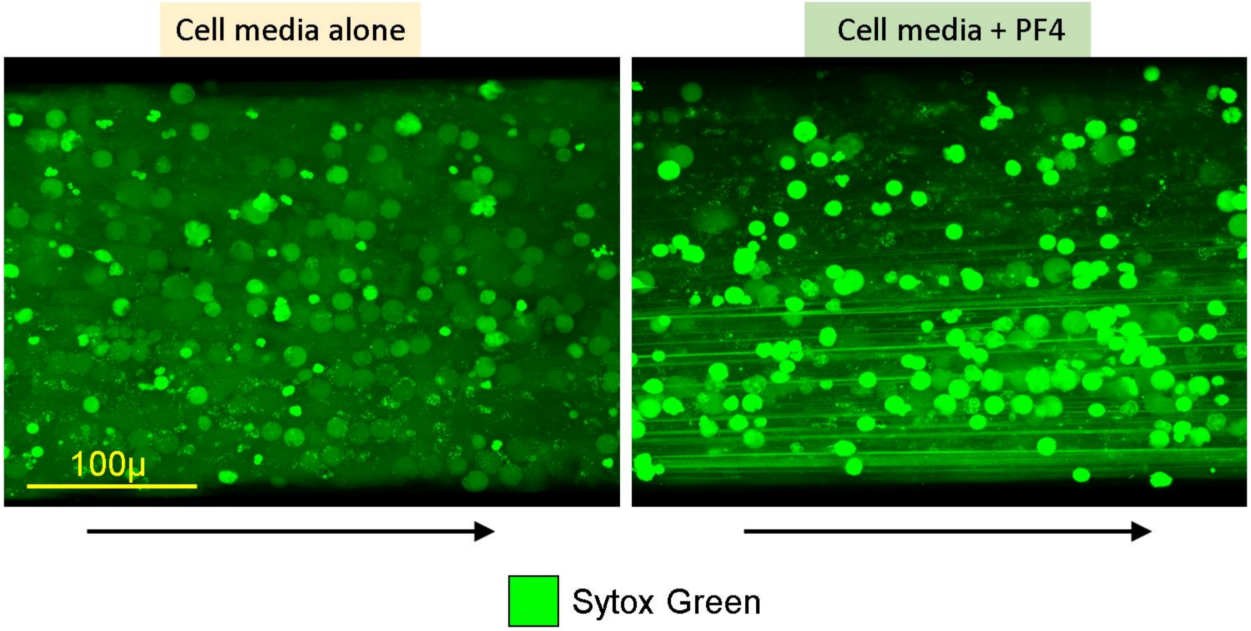


**Supplement Figure 1. cfDNA released from neutrophils adherent to HUVECs in a microfluidic chamber.**

Representative fields of released cfDNA from LPS-stimulated neutrophils in a HUVEC lined microfluidic channel demonstrating the compaction and aggregation of the cfDNA by the addition of hPF4 (25 µg/ml). Arrow indicates direction of flow. Size bar included as well. Images obtained with a Zeiss LSM 710 confocal microscope. Original magnification was 20x.

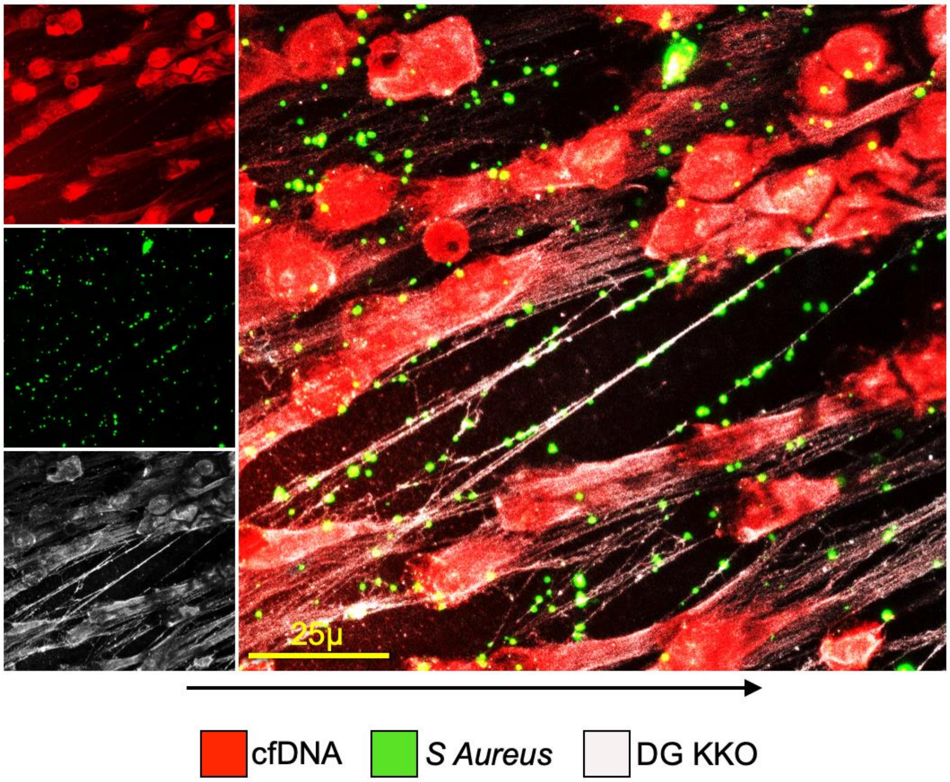


**Supplement Figure 2. Entrapment of S aureus bacteria on DG-KKO/PF4-NET complexes in a microfluidic well.**

Representative confocal image of SYTOX orange-stained NETs compacted with PF4 (100 µg/ml) and bound by 647-labeled DG-KKO, infused with 488-labeled *S Aureus* bioparticles (40 µg/ml), showing direct adhesion of *S aureus* bioparticle to DG-KKO complexed to PF4-NETs. Images obtained with a Zeiss LSM 710 confocal microscope. Original magnification 40x. Scale bar and arrow showing direction of flow are indicated.

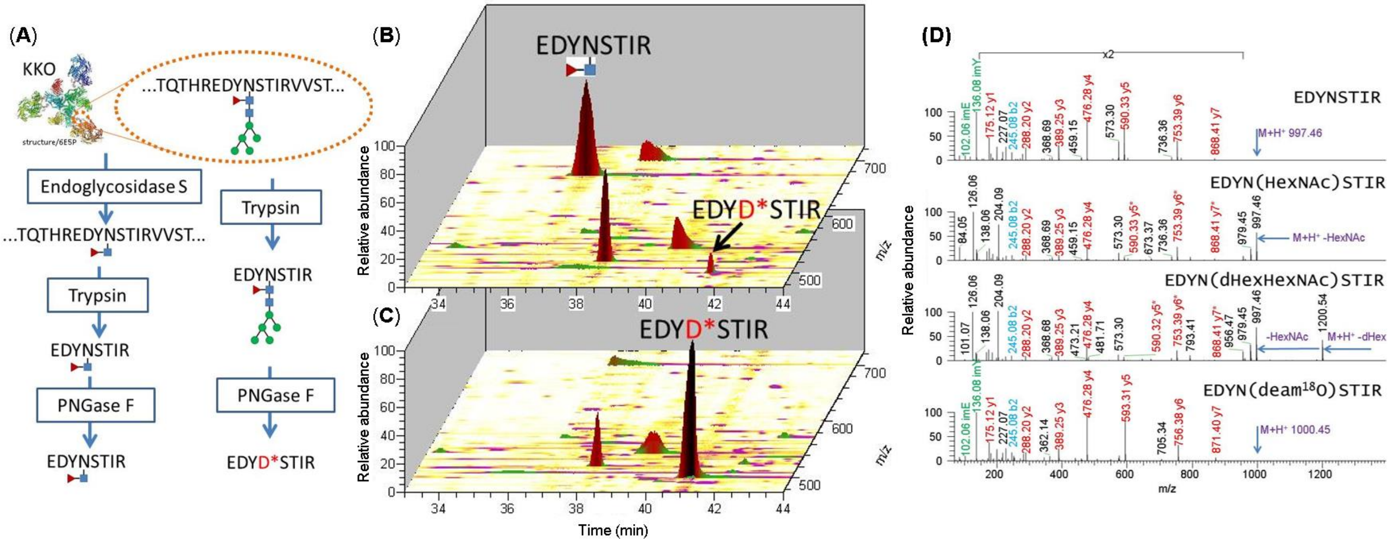


**Supplement Figure 3. MS/HPLC analysis confirms the deglycosylation of KKO.**

(A) Schematic describing DG-KKO generation by incubation with endoglycosidase S and preparation for analysis with mass spectroscopy with treatment with trypsin and PNGase F (on left) compared to control KKO. (B) Analysis of peptide fragments generated following antibody incubation with endoS showing a large peak at 37 minutes representing 93% of the sample corresponding to a fucosylated peptide generated following digestion of DG-KKO, and a small peak at 42 minutes representing the deiminated peptide generated from KKO. (C) Analysis of peptide fragments generation from digestion of KKO, showing no peak at 37 minute, but a large peak at 42 minutes corresponding to the deiminated peptide. (D) Amino acid sequences of peptides generated from the digestion of KKO and DG-KKO. The top two sequences cumulatively represent 3% of the sample. The fucosylated peptide (3rd from the top) was 93% of the peptides obtained from digestion of DG-KKO. The deminiated peptide (4th from the top) represented ~6% of the peptides generated from DG-KKO and 100% of the peptides generated from KKO.

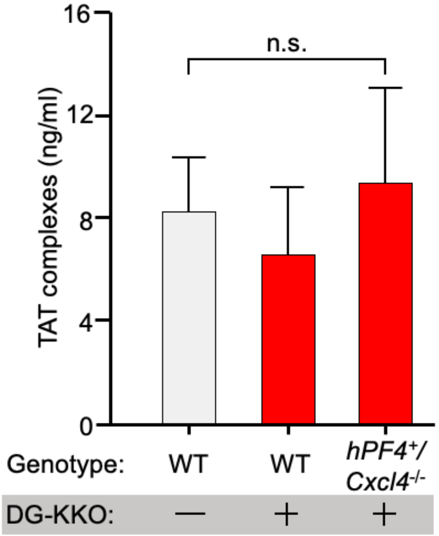


**Supplement Figure 4. TAT levels in DG-KKO-treated LPS-exposed mice.**

Mice were injected with LPS (35 mg/kg, IP). 30 minutes later they received tail vein injections containing vehicle alone or 5 mg/kg DG-KKO. 6 hours following LPS injection, mice were euthanized and IVC blood samples were collected and plasma was isolated. ELISA was used to quantify TAT complexes concentrations in the plasma samples. N = 3-9 animals per arm. n.s. = no significant difference. Statistical analysis was performed with a Kruskal-Wallis one-way ANOVA.

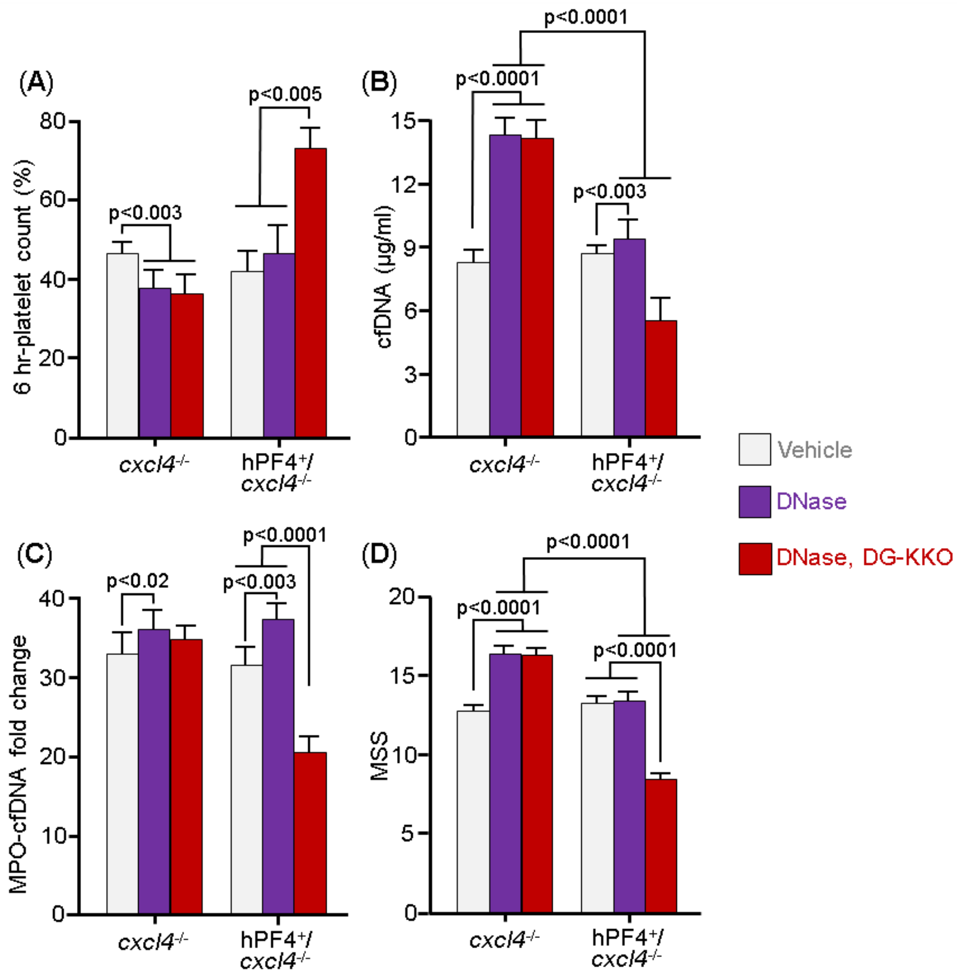


**Supplement Fig 5. DG-KKO effects on platelet count, NDP release and survival in LPS ± DNase I-exposed hPF4$^{+}$ mice.**

Mice received LPS (35 mg/kg body weight, IP) and DNase I (20 mg/kg, IP). Mean ± 1 SD of (**A**) platelet counts, (**B**) cfDNA, (**C**) MPO-cfDNA, and (**D**) MSS were measured at 6 hours post-exposure. N=10 animals per arm. Analysis performed with Sidak's multiple comparison t-test.
